# Semi-supervised segmentation and genome annotation

**DOI:** 10.1101/2020.01.30.926923

**Authors:** Rachel C.W. Chan, Matthew McNeil, Eric G. Roberts, Mickaël Mendez, Maxwell W. Libbrecht, Michael M. Hoffman

**Affiliations:** University of Toronto; Princess Margaret Cancer Centre; Simon Fraser University; Vector Institute

## Abstract

Segmentation and genome annotation methods automatically discover joint signal patterns in whole genome datasets. Previously, researchers trained these algorithms in a fully unsupervised way, with no prior knowledge of the functions of particular regions. Adding information provided by expert-created annotations to supervise training could improve the annotations created by these methods. We implemented semi-supervised learning using virtual evidence in the annotation method Segway. Additionally, we defined a positionally tolerant precision and recall metric for scoring genome annotations based on the proximity of each annotation feature to the truth set. We demonstrate semi-supervised Segway’s ability to learn patterns corresponding to provided transcription start sites on a specified supervision label, and subsequently recover other transcription start sites in unseen data on the same supervision label.

## 1 Introduction

The rapid evolution of high-throughput DNA sequencing in the past decade has resulted in a huge number of whole-genome datasets. In multitudes of samples and tissue-types, there exist datasets which measure histone modification, DNA methylation, transcription factor binding, and other physical and chemical properties. To make sense of data at this scale, genomics researchers use computational methods to identify patterns which shed light on the underlying genomic activity.

For this purpose, a class of methods known as segmentation and genome annotation (SAGA) methods have become popular. The Segway^1,2^ and ChromHMM^3,4^ methods provide two examples of such methods. Most SAGA methods, including ChromHMM, only learn in an unsupervised mode, meaning they discover genomic signal patterns without any prior information about known categories of genomic elements. Likewise, currently researchers use Segway primarily as an unsupervised annotation method.

For many use cases, one could guide a SAGA method to better performance by providing supervision information during training. For example, we have high-quality data on experimentally verified genes, promoters, and enhancers. This information might prove useful by influencing the patterns Segway learns in these cases, or by directing it to find other regions of the same pattern. This semi-supervised learning task differs from both unsupervised and fully supervised learning in that we provide both labeled and unlabeled training data to the model.

We implemented semi-supervised learning in Segway’s graphical model using virtual evidence. Virtual evidence provides a way of specifying prior information (“priors”) about a set of probabilistic events in a Bayesian model^5^ by encoding the prior information into binary dummy observed nodes. One specifies that the probability of the dummy node given some explanation equals the probability of the evidence for the explanation. For example, we might have a prior belief in a hypothesis that a specific genomic position has a particular behavior, or chromatin state. Using virtual evidence, we can include in the model our belief in the position’s behavior. To do this, we specify a probability representing the credibility of our evidence towards our hypothesis. When we supply virtual evidence for at least some parts of the training data, virtual evidence provides a framework for semi-supervised learning.

## 2 Methods

### 2.1 Transcription start site prediction

Detection of transcription start sites (TSSs) in the genome is a well-studied problem in computational genomics.^6–8^ Accurate prediction of TSSs and promoters can guide the discovery and verification of previously-unknown genes. It can also identify candidate genomic regions for further experimental search.^9^

Toward the goal of accurate genome and TSS annotation, researchers have developed many algorithms to predict TSSs from genomic sequence^10–14^ and to identify genes from RNA-seq data.^15–17^ No existing methods, however, predict TSSs on histone modification and DNase data alone. Histone modifications and DNase have characteristic patterns of signal in the regions surrounding distinct regulatory elements, such as surrounding TSS.^18,19^ cap-analysis gene expression (CAGE) provides information on the 5′ ends of capped RNA transcripts.^20^ This distinguishes it from RNA-seq, which only provides random tags along the length of steady-state expressed RNA.^21^ As such, RNA-seq only provides information on steady-state transcript expression instead of transcription initiation. RNA-seq therefore has limited application in the study of TSS regulation.

We used FANTOM5 CAGE data in conjunction with histone modification and DNase data in cell types with all three types of data available. Given that CAGE provides information on the specific locations of TSSs, this approach could lead to better understanding the relationship between histone modifications and transcription initiation. We trained on histone modification and DNase data while using CAGE data as supervision labels, and predicted the locations of TSSs on only histone modification and DNase data. We leveraged an existing TSS reference database, intersected with CAGE data, to create a cell type-specific TSS set for derivation of supervision labels and for scoring the model on test data. We created methods to evaluate TSS prediction within the context of uncertainty in determining which TSSs are active in some cell type.

Despite some uncertainty in determining which promoters have activity in specific cell types, promoters remain one of the most straightforward genomic regulatory elements to validate. For this task, we can leverage multiple comprehensive databases of gene features.^22–25^ In comparison, enhancer prediction has proven considerably more difficult.^26–28^ This difficulty arises, in part, from enhancers’ location distal from the genes whose transcription they modify.^29–33^ In addition, unlike promoters, researchers have not yet created comprehensive catalogues of enhancers—there likely exist many yet to be discovered.

We aim to develop a predictive method that can be used for predicting many different types of genomic elements. Promoter prediction makes an excellent baseline task for the development of this method. One could then apply it to similar tasks with less straightforward performance evaluation, such as including enhancers and insulators.^34^

### 2.2 Segway’s graphical model

Segway uses a dynamic Bayesian network which models a segment label at each genomic position *t* ∈ {0, … , *T*} as a hidden discrete random variable *Q*_*t*_ ∈ {0, … , *ζ*}, and models signal datasets *i* ∈ {0, … , *I*} as observed continuous random variables 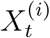. For each (position, dataset) pair (*t, i*), Segway also has an indicator “presence” random variable *X̊*_*t*_^(*i*)^ to account for missing signal data.

The value of the presence variable affects the strength of the relationship between *Q*_*t*_ and 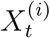 through a conditional dependence structure. Specifically, the value of *X̊*_*t*_^(*i*)^ determines the conditional dependence relationship between *Q*_*t*_ and 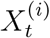 by the following rules: 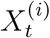 and *Q*_*t*_ are conditionally independent if *X̊*_*t*_^(*i*)^ = 0, and not conditionally independent otherwise. Using notation from Lauritzen [35], we denote two conditionally independent random variables □ and ◯ given some condition △ as □ ⫫ ◯ | △. Correspondingly, we can express the presence variable conditional independence rules as

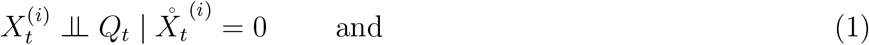

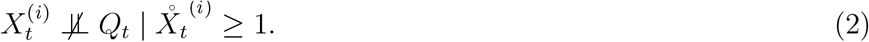

For example, if position *t* has missing data from signal dataset *i*, then *X̊*_*t*_^(*i*)^ = 0, which implies no dependence between *Q* and 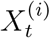. In other words, *X̊*_*t*_^(*i*)^ = 0 means 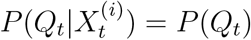. As a result, the information at 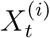 is not used by the rest of the model.

The presence structure proves particularly useful when running Segway at a resolution coarser than 1 bp. The structure takes into account how many positions in a resolution-length window have data defined, and weights the contribution of the resulting downsampled window accordingly by replacing *P*(*X*_*t*_^(*i*)^) in the likelihood function with 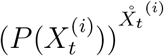. This is a generalization of the behavior at resolution 1, where *X̊*_*t*_^(*i*)^ takes on values 0 or 1.

To represent different behaviors using labels, Segway learns mixture models with *K* Gaussian components. Variance parameters are tied across signal datasets, in the sense that all labels share the same variances for a particular dataset. For each (dataset, label) pair (*i, q*), this results in *K* mean parameters *μ*_*ikq*_. But for each dataset *i*, all labels share the same *K* variance parameters 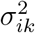. When *K* = 1, Segway uses a simple Gaussian model with one Gaussian per label.

During training, Segway uses expectation-maximization (EM)^36^ to estimate parameters of its mixture models. During the subsequent annotation task, Segway partitions the genome into non-overlapping segments, and uses the labels learned to assign one label to each resulting segment. Segway assigns labels such that regions with the same label share similar properties in the observed data and may correspond to functions such as “promoter” or “enhancer”. Segway can learn with EM over the entirety of specified training regions, or use minibatch training on a random fixed percentage thereof.

### 2.3 Virtual evidence in Segway

#### Graphical model structure

To add semi-supervised learning into Segway, we augmented its graphical model with virtual evidence. Specifically, we added an observed binary virtual evidence node *V*_*t*_ at each position *t* which has the hidden segment label *Q*_*t*_ as its parent (Figure 1), and always has value unity, *V*_*t*_ ≡ 1. The virtual evidence node specifies our prior belief in the value of *Q*_*t*_ given only some uncertain evidence. It is a dummy node that propagates the uncertainty *ɛ* in evidence for some value of *Q*_*t*_, represented by *P*(evidence explanation) = *P*(*V*_*t*_ = 1|*Q*_*t*_ = value) = *ɛ*. For example, specifying *P*(*V*_*t*_ = 1|*Q*_*t*_ = 2) = 0.3 means that we have uncertain evidence that *Q*_*t*_ has a value of 2 with probability 0.30. Since *V*_*t*_ = 1, we can use Bayes’ theorem to compute the posterior probability *P*(explanation|evidence) ∝ *P*(evidence|explanation)*P* (explanation).^37^

**Figure 1:**
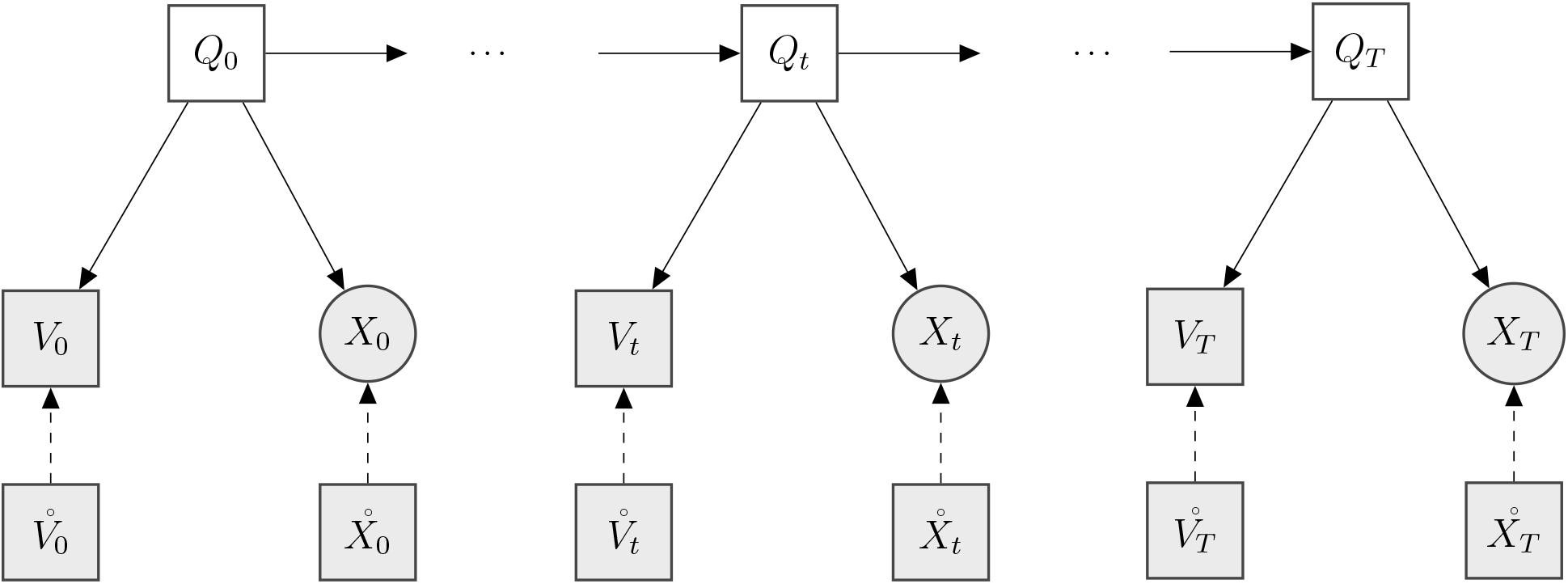
An abridged model of Segway with virtual evidence. The binary virtual evidence child *V*_*t*_ specifies the uncertainty in evidence for its parent *Q*_*t*_ taking on a particular value *q*_*t*_ by defining the conditional probability vector *P* (*V*_*t*_ = 1|*Q*_*t*_ = *q*_*t*_). As with the signal variable *X*_*t*_ and associated presence variable *X̊*_*t*_, the influence of *V*_*t*_ is deterministically controlled by the associated presence variable *V̊*_*t*_. The value of *V̊*_*t*_ is the number of positions with priors specified in its corresponding resolution-length window. Node shape: discrete (square) or continuous (circle). Node color: hidden (white) or observed (grey). Solid arcs represent conditional dependence between random variables; dashed arcs indicate that the child is deterministically weighted by the parent.

Our virtual evidence graphical model structure allows us to specify a vector of priors at each position *t*. This vector has one value for each realization *Q*_*t*_ = *q*_*t*_, where *q*_*t*_ ∈ {0, … , *ζ*}. The vector corresponds to the probability that *V*_*t*_ = 1 given each realization of *Q*_*t*_, *P*(*V*_*t*_ = 1|*Q*_*t*_ = *q*_*t*_). And we defined *V*_*t*_ to have the realization 1 always.

With no specified priors for a particular position, the vector of used priors defaults to the uniform priors *P*(*V*_*t*_ = 1|*Q*_*t*_ = *q*_*t*_) = 1/(*ζ* + 1). With partially specified priors, we infer uniform probability for the remaining *q*_*t*_ values. For example, if we only have pre-existing beliefs for *Q*_*t*_ = 0 with *ζ* = 3, then 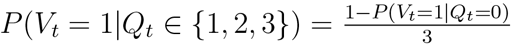.

#### Downsampling virtual evidence to lower resolution

To perform inference at a resolution coarser than 1 bp, we partition the genome into resolution-length windows. We then downsample virtual evidence by averaging the priors of positions which have specified priors in each resolution-length window. Using a similar presence structure as described for 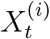, we define the discrete observed presence variable *V̊*_*t*_^(*i*)^ which counts the number of positions in a resolution-length window which have specified priors. In an identical manner to *X̊*_*t*_^(*i*)^, *V̊*_*t*_^(*i*)^ weights the strength of the averaged priors’ contribution to the graphical model by replacing *P*(*V*_*t*_) in the likelihood function with 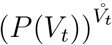.

### 2.4 Experimental setup

To demonstrate the utility of virtual evidence in Segway, we applied it predicting TSSs in the leukemia cell line K562^38^ (Figure 2a). We trained Segway on ENCODE Project Consortium^39^ GRCh38^40^/hg38 DNase-seq data and ChIP-seq datasets for H3K27ac, H3K4me3, H3K36me3, and H3K27me3 (Table S1). With H3K27me3 and H3K36me3, we generally expect depletion at TSSs whereas with H3K4me3 and H3K27ac, we generally expect enrichment at active TSSs.^41–45^ We used a 3-component Gaussian mixture model for each label to enable Segway to learn more complex signal distributions.

**Figure 2:**
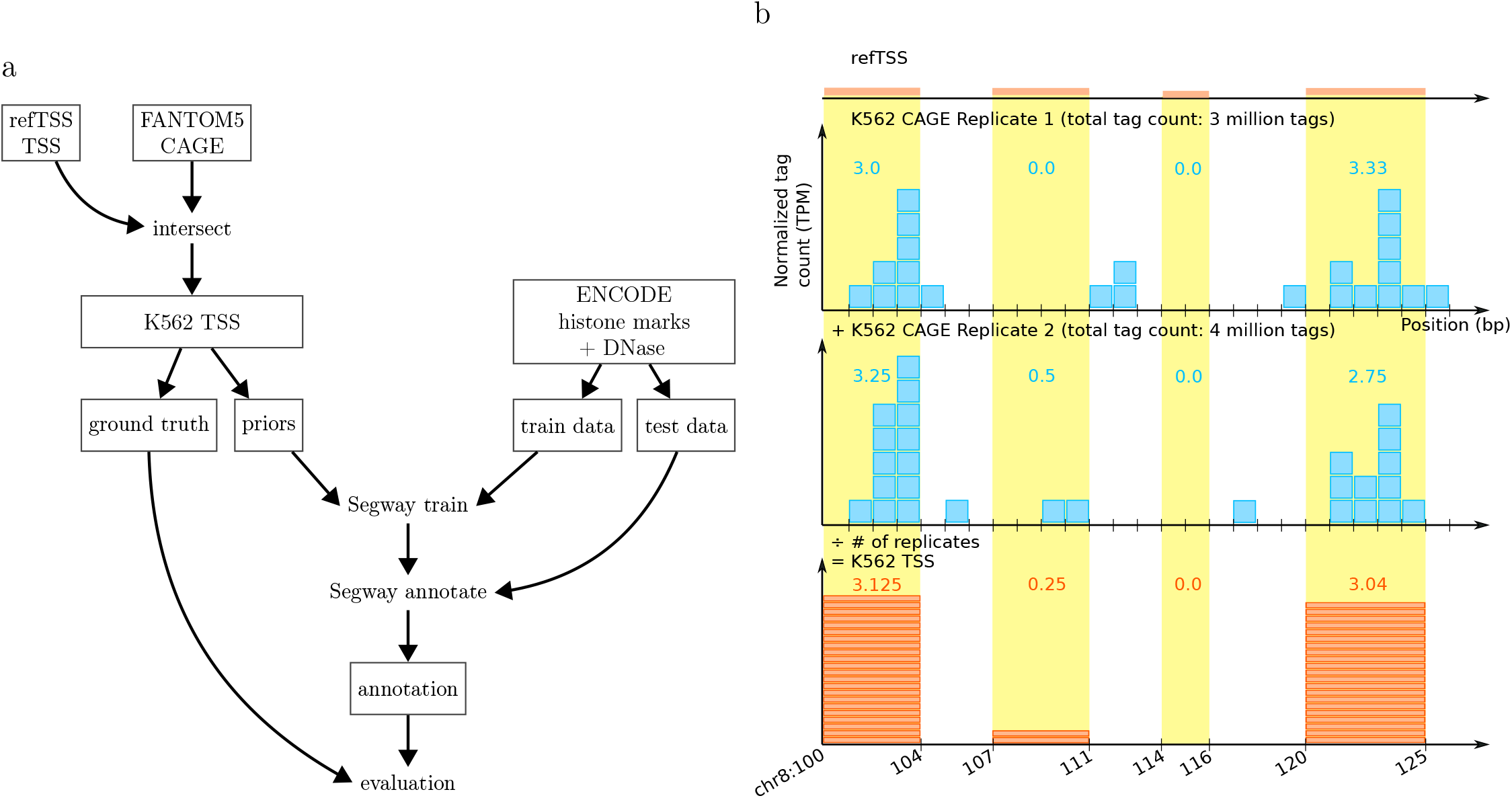
Experimental workflow and intersection of refTSS d atabase with K562 CAGE replicates. **(a)** Experimental workflow, s tarting from genomic databases a nd e nding w ith evaluation of Segway annotation. Boxed nodes indicate data objects. Boxed nodes with multiple rows of text: 1st row indicates data source; other rows describe data. Unboxed nodes indicate actions performed upon the incoming data nodes, resulting in the outgoing nodes. **(b)** Toy example illustrating intersection of refTSS database with 2 CAGE replicates to create K562 TSS. First, we normalize the CAGE tag count of each replicate by converting its tag counts to TPM. Next, for each refTSS feature with support from either replicate, we sum the normalized tag counts of the replicate features overlapping it, then divide this value by the total number of replicates. This creates a K562-specific s ubset of refTSS (“K562 TSS”) with corresponding averaged normalized tag count information.

We ran Segway with and without virtual evidence, and compared the performance of each case. In both cases, we used a resolution of 1 bp, a total of 10 labels with 10 random starts, and trained for 40 EM iterations. We used the same random seed for both the unsupervised control and Segway with virtual evidence.

To account for the cell-type–specific nature of transcription, we combined information from K562 CAGE with a cell-type–agnostic TSS database to form a K562-specific s et of TSS (“K562 T SS”). For a CAGE experiment with some total number of tags sequenced (“total tag count”), De Hoon et al. [46] define the tags per million (TPM) of a feature as

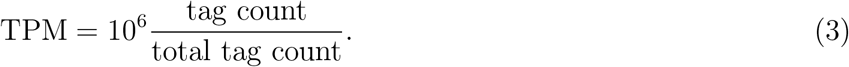

To create a large window around each TSS on which to train, we increased the size of + strand K562 TSS^47^ features with high CAGE support (TPM ≥ 2.5) by 10 000 bp on both sides. We trained Segway on these expanded regions on chromosome 21 (1.0% of the total set of expanded regions across all chromosomes) and annotated on all chromosomes.

We chose to train only on + strand K562 TSS. ChIP-seq and DNase-seq do not report properties which are strand-specific. Promoters, however, may have distinct histone modification patterns upstream and downstream.^48^ Training on promoters from both strands would require a more complex transition model that could switch transition structures depending on the direction of the promoter. As a first attempt at promoter prediction, we avoided this complexity by training only on the + strand.

To evaluate Segway’s performance, we defined positionally tolerant precision and recall metrics for scoring genome annotations based on the proximity of each annotation feature to the truth set. We evaluated each of unsupervised Segway and Segway with virtual evidence using positionally tolerant precision and recall on the truth set. To do this, we computed positionally tolerant precision and recall based on Segway’s annotation performance on chromosomes 1–20, 22, and X.

#### Ground truth and priors

We formed our ground truth and prior values by combining CAGE data^22,23,49,50^ with the refTSS database.^47^ CAGE captures and extracts the 5′ ends of capped RNA transcripts. Thus, its output fragments, known as CAGE tags, correspond to transcript starts. We call the 5′ end of an aligned CAGE read a “ CAGE tag starting site”.

The refTSS database contains human transcription start sites, formed by integrating multiple TSS annotations and resources, such as FANTOM5, DBTSS,^51^ ENCODE CAGE,^52^ and ENCODE RAMPAGE.^53^ First, we normalized the CAGE tag counts of all biological replicates available in K562 by converting the tag counts of each replicate to TPM. Second, to get cell-type–specific TSSs from the cell-type–agnostic refTSS, we overlapped refTSS with the K562 CAGE tag starting sites of each of these biological replicates. For the resulting K562 TSS features, we summed the overlapping normalized tag counts from the overlapping features (Figure 2b). Third, we then averaged the summed normalized tag counts of all K562 TSS features by dividing by the total number of replicates. To compute our priors, we removed K562 TSS features with TPM < 2.5. As 74.2% of K562 TSS features have TPM < 2.5, this creates a highly stringent set of high-confidence active TSSs.

For every K562 TSS feature, we derived a prior probability of transcriptional activity from an affine transformation of the feature’s expression level. The base confidence h yperparameter *β*_0_ expresses our confidence in the whole set of K562 TSS features with TPM ≥ 2.5. The relative confidence hyperparameter *β*_1_ expresses how much our confidence increases with increased expression, measured in TPM per feature base pair. For any K562 TSS feature with TPM ≥ 2.5 and length *ℓ*, we computed its prior probability as

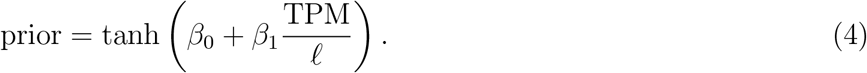

Dividing by feature length indicates that we have more confidence in a small feature with a high TPM, than a large feature with the same TPM. Finally, we apply a hyperbolic tangent transformation. Since all inputs are positive, this ensures that the resulting value’s range varies between 0 and 1.

To form our ground truth, we filtered the K562 TSS set and kept only features with TPM ≥ 0.5. We used a threshold of TPM ≥ 0.5 as 49.3% of K562 TSS features have TPM < 0.5. In comparison to using a threshold of TPM ≥ 2.5, this created a stringent set of active TSSs, yet still retains at least half of all K562 TSS to score against.

Here, we used base confidence *β*_0_ = 1.50 and relative confidence *β*_1_ = 0.02 bp/TPM. We chose our base confidence to encapsulate the intuition that features with TPM at least 2.5 are highly likely active TSSs, since tanh(1.50) ≈ 0.90. We chose our relative confidence because 99.6% of those features had length-normalized TPM values above the threshold of 50 TPM/bp. We specified a relative confidence of the reciprocal of this threshold, 1/(50 TPM/bp) = 0.02 bp/TPM.

We expanded by 200 bp on both sides each + strand K562 TSS feature with TPM ≥ 2.5. We merged any expanded TSS features which overlapped, setting the merged region’s prior to the average of the overlapping features. We used these expanded and merged regions as supervision regions, with the arbitrarily chosen label 0 as our supervision label.

#### Positionally tolerant precision and recall metrics for genomic predictions

To evaluate our method, we developed positionally tolerant metrics similar to precision and recall. These metrics measure an annotation’s performance once in an environment where we do not need (or cannot obtain) base pair accuracy. Biologists using annotations from SAGA methods do not need complete positional exactness when investigating some labeled region. Moreover, many epigenomic assays, including ChIP-seq, provide only inexact information on the location of regulatory elements. To account for these circumstances, we define positionally tolerant precision and recall, which one can apply to compare any predicted annotation with a ground truth annotation.

These metrics are not true precision and recall metrics as they use different numerators from each other. Instead, the positionally tolerant metrics provide a way of quantifying performance that satisfies much of the intuition that one might get from standard precision and recall. Since the positionally tolerant metrics are not true precision and recall metrics, various invariants with respect to conventional precision and recall may not hold in these definitions.^54,55^

The positionally tolerant metrics summarize extended overlap between two sets of closed, non-overlapping intervals: a prediction set *Ŷ* and a ground truth set *Y*. They arise from a number of comparisons between a single prediction interval *ŷ* = [*a*_1_, *a*_2_], and a single ground truth interval *y* = [*b*_1_, *b*_2_], with *a*_1_, *a*_2_, *b*_1_, *b*_2_ ∈ ℕ^0^. We define the distance between these intervals *d*(*ŷ*, *y*) to be the smallest distance between any point in each interval to any point in the other,

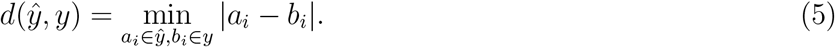

For a single closed interval *ŷ* = [*a*_1_, *a*_2_] and the whole ground truth set *Y*, we define the interval–to– interval-set distance as

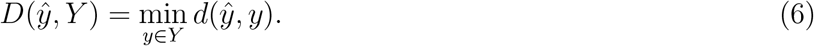

*D*(*ŷ*, *Y*) gives the shortest absolute distance from interval *ŷ* to any interval in the set *Y*.

To represent a tolerance for considering an interval proximate to an interval set, we define a radius *r* ∈ ℕ^0^. We can then define positionally tolerant precision and recall by computing the fraction of intervals in each set which are proximate to the other set. With radius *r*, the positionally tolerant precision P_*r*_ between these sets is then

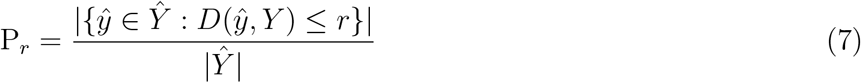

and the positionally tolerant recall R_*r*_ is

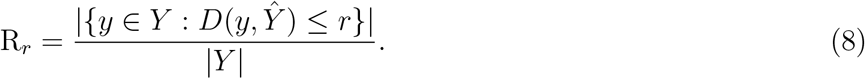

When *r* = 0 and we decompose each interval into a sequence of contiguous intervals, each with length 1 bp, positionally tolerant precision and recall become equivalent to non-tolerant pointwise precision and recall.

P_*r*_ represents the fraction of prediction segments which have interval–to–interval-set distance to the ground truth less than or equal to *r*. Similarly, R_*r*_ represents the fraction of ground truth segments which have interval–to–interval-set distance to the prediction less than or equal to *r*.

Applied to the TSS prediction problem (Figure 3), we can rewrite the formulas as

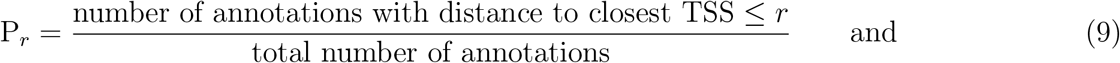

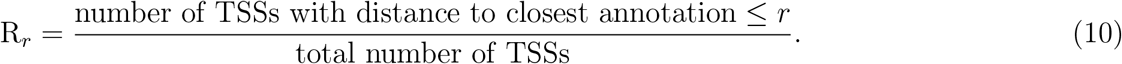

**Figure 3:**
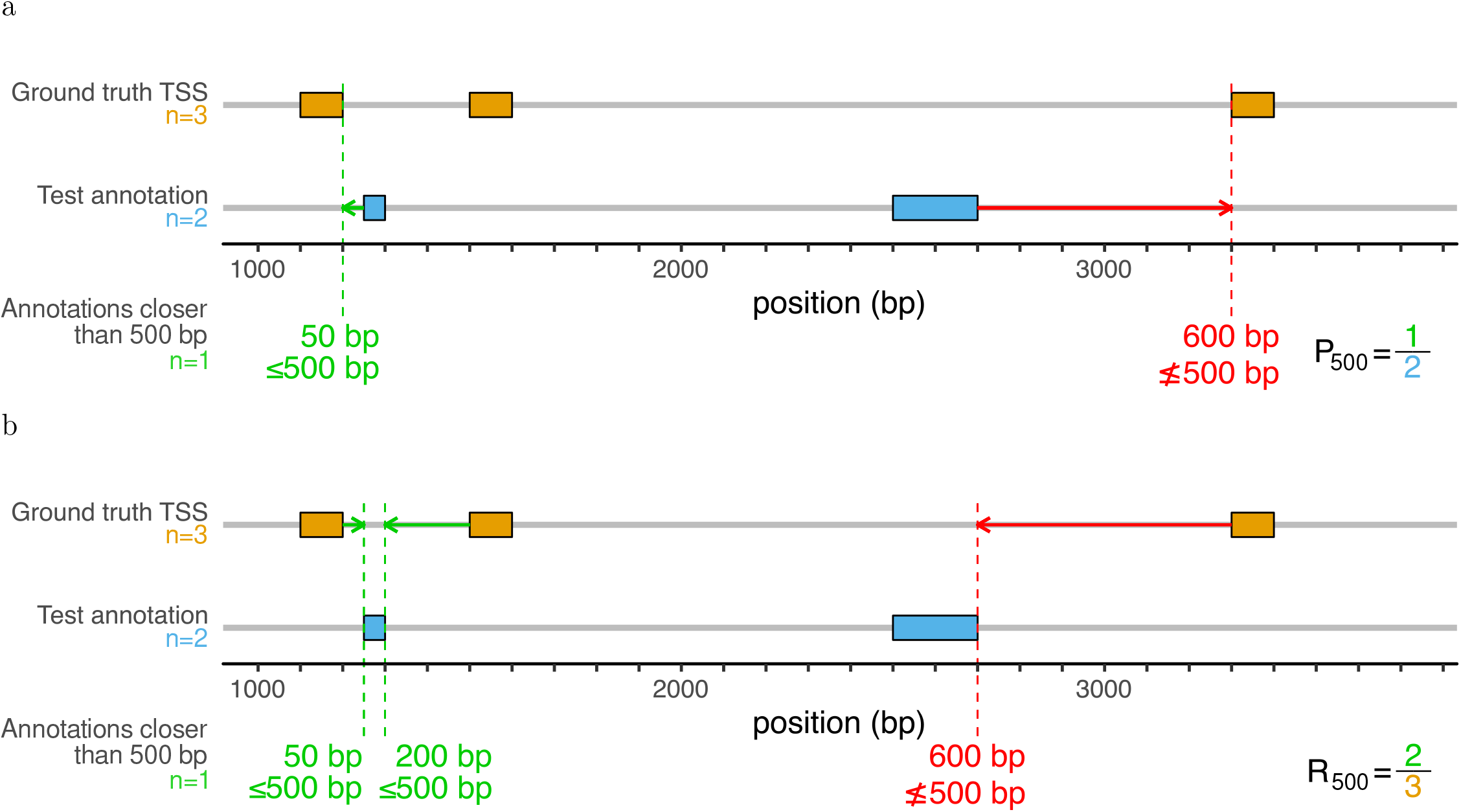
Toy example demonstrating positionally tolerant precision and recall with a radius of 500 bp. **(a)** Computation of positionally tolerant precision. The test annotation has 2 features, so we perform 2 comparisons to compute the positionally tolerant precision. To compute the positionally tolerant precision, we first find the number of an notations wi th di stance to closest TSS le ss th an or eq ual to 500 bp. The 2 annotations have closest distances 50 bp and 600 bp. Only 1 of these distances is less than or equal to 500 bp, so the positionally tolerant precision is accordingly 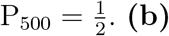. **(b)** Computation of positionally tolerant recall on the same test annotation and ground truth TSS set as (a). The ground truth TSS set has features, resulting in 3 comparisons. This is 1 more than for the positionally tolerant precision computation, since the test annotation only has 2 features. To compute the positionally tolerant recall, we find the number of TSSs with distance to closest annotation less than or equal to 500 bp. The 3 annotations have closest distances 50 bp, 200 bp, and 600 bp. Since 2 of these distances are less than or equal to 500 bp, the positionally tolerant recall is accordingly 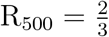.

We can choose the radius *r* to increase or decrease positional specificity. Histone modification ChIP-seq signals surrounding TSSs are generally enriched at a scale of 2000 bp or greater.^42,56–58^ We chose a radius of 500 bp here to allow for positional specificity on a comparable resolution while preventing excessive permissiveness in classification.

## 3 Results

### 3.1 Virtual evidence improved performance relative to an unsupervised model

We assessed Segway’s ability to predict TSSs on either strand both with and without virtual evidence during training (Table 1). We used our ground truth of K562 TSS features with TPM ≥ 0.5 located on all chromosomes except training chromosome 21. As expected, virtual evidence caused the supervision label (label 0) to capture the most TSSs. In comparison, the unsupervised control happened to place most TSSs on label 3. We considered label 3 the top-performing label for the unsupervised control, as it had a higher harmonic mean of positionally tolerant precision and recall (0.55), compared to the next top-performing label, label 7 (harmonic mean 0.47). Virtual evidence’s placement of most TSSs on a different label from the unsupervised control demonstrated that virtual evidence successfully influences the labels that Segway discovers.

**Table 1:** Comparison of two modes of identifying TSSs: virtual evidence and an unsupervised control.

|  | Supervision label<sup>a</sup> | Evaluated label<sup>b</sup> | Precision<sup>c</sup> P<sub>500</sub> | Recall<sup>c</sup> R<sub>500</sub> |
| --- | --- | --- | --- | --- |
| Unsupervised | None | 3 | **0.44** | **0.72** |
|  |  | 7 | 0.33 | 0.81 |
| Virtual evidence | 0 | 0 | **0.69** | **0.76** |

<sup>a</sup> Label on which TSSs were supervised, if applicable.
<sup>b</sup> Label with the highest precision and recall in identifying TSSs on unseen data. For the unsupervised control, we reported the top 2 performing labels.
<sup>c</sup> Positionally tolerant, with radius 500 bp. Test set: the human genome except chromosome 21. Bold: best performance for the mode.

Virtual evidence directs particular labels towards patterns found in the supervision region. It then finds other, similar regions in data unseen at training. Using virtual evidence, Segway obtained a 24% increase in precision and 4% increase in recall from the unsupervised control (Table 1).

To understand the comparative performance between Segway with virtual evidence and its unsupervised control, we examined a region around the *HMBS* gene^59^ (Figure 4). *HMBS* expresses a protein involved in the synthesis of heme. As K562’s biological source is human blood, we expect to see transcriptional activity at this gene. At the *HMBS* TSS, we found that both Segway with virtual evidence and its unsupervised control were able to correctly identify the region as a TSS.

**Figure 4:**
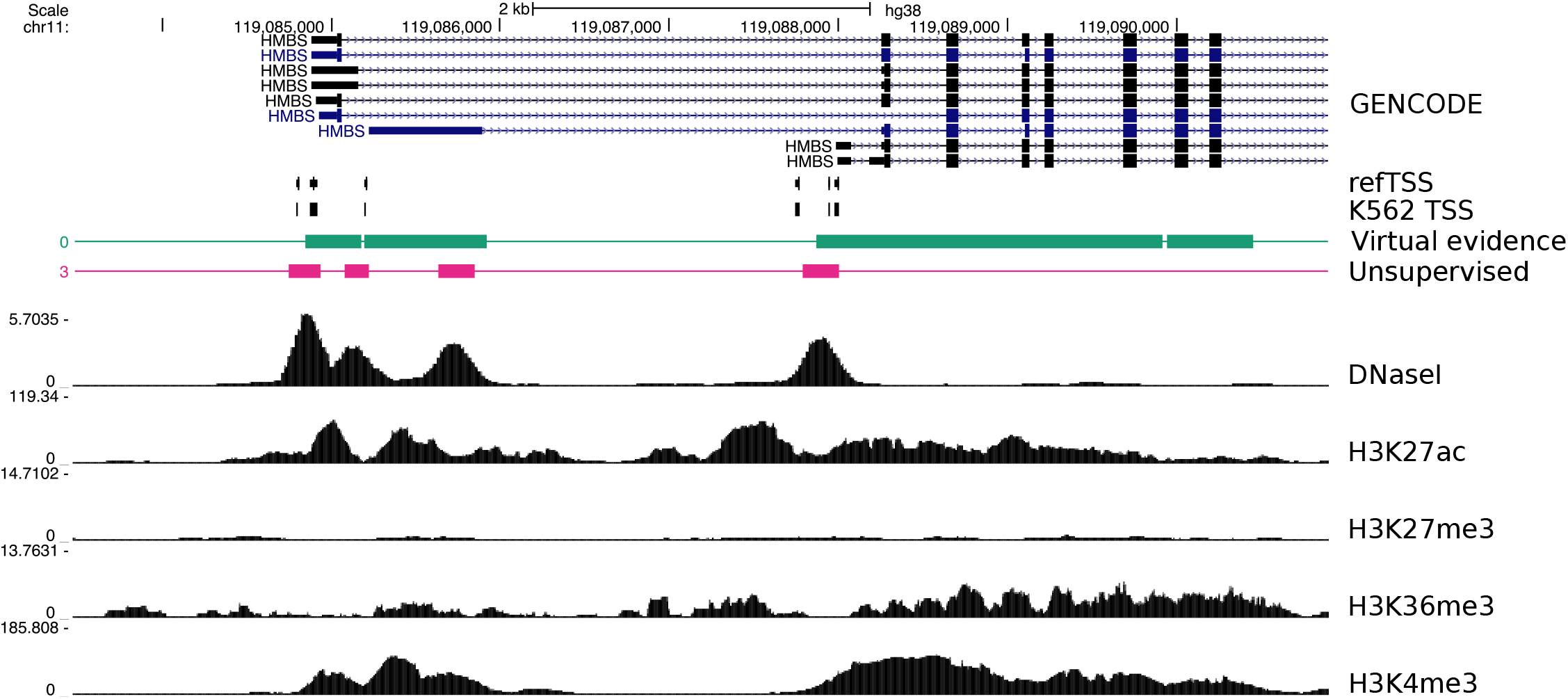
UCSC Genome Browser^60^ display of the *HMBS* locus. Both virtual evidence (supervised label 0) and the unsupervised control (label 3) captured the K562 TSS peak. We show the basic gene annotation set from GENCODE^25^ version 32. Below it, we show the refTSS database and cross-referenced K562 TSS set. Finally, we show the corresponding signal data for DNase-seq and H3K27ac, H3K27me3, H3K36me3, and H3K4me3 ChIP-seq (Table S1). The maximum viewable value for each signal track is its 99.95th percentile value across the whole genome, the minimum range which permits visualization of all signal data without clipping.

Additionally, we examined the region around the *DAB1* gene (Figure 5). The unsupervised control incorrectly identified a TSS within the *DAB1* gene, but not at the *DAB1* TSS. As expected, this non-TSS region has no refTSS or GENCODE support. In comparison, Segway with virtual evidence did not identify this region as a TSS, contributing to an increase in precision compared to the unsupervised control.

**Figure 5:**
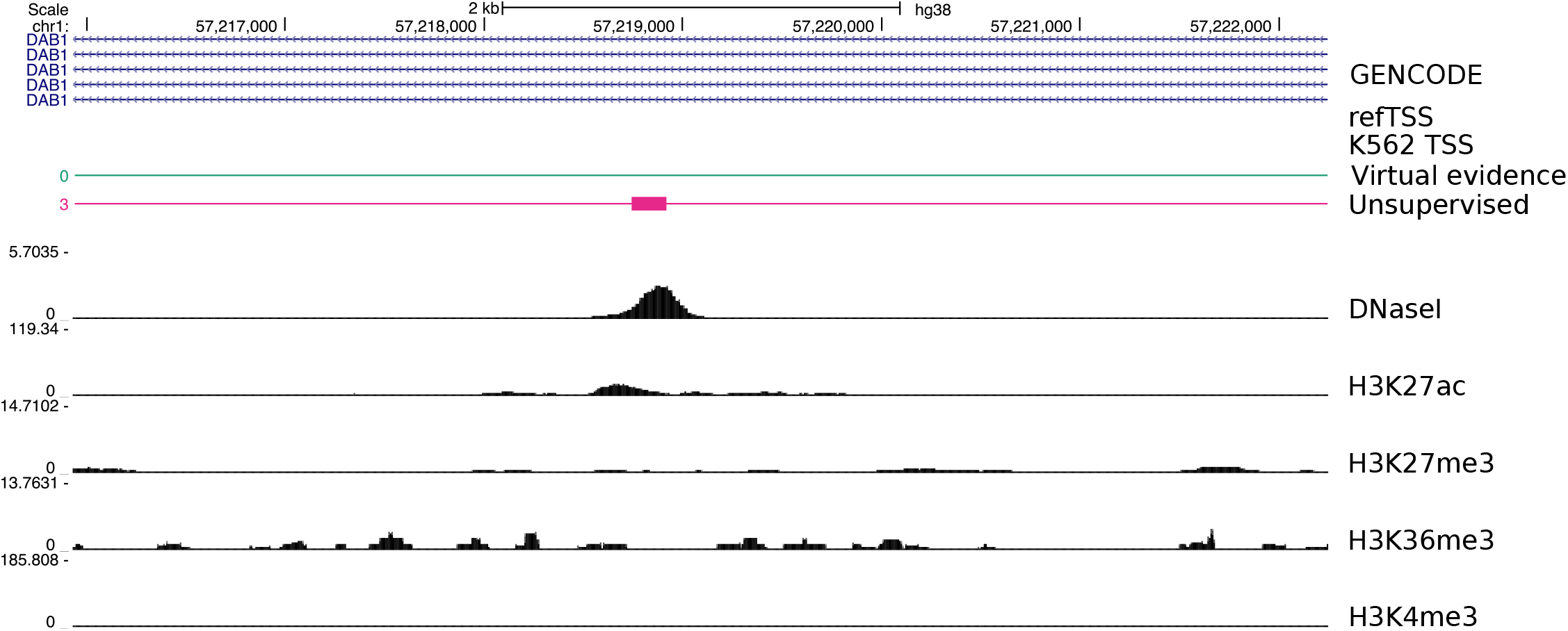
UCSC Genome Browser^60^ display of the *DAB1* locus. The unsupervised control (label 3) incorrectly labels as TSS a region with no refTSS or GENCODE support. In comparison, virtual evidence (supervised label 0) does not label this region as TSS, contributing to an increase in precision (Table 1). Meaning of tracks is identical to Figure 4.

As a baseline for evaluation, we used a positionally tolerant precision and recall radius of 500 bp, chosen based on the patterns of enrichment of histone modification ChIP-seq signal surrounding TSSs. To better understand the effect of varying the radius, we also computed the positionally tolerant precision and recall using radii ranging from 1 bp to 500 bp (Table S2). Increasing the radius from 1 bp to 50 bp led to a disproportionate gain in recall, compared to the relatively incremental increase obtained by going from 50 bp to 100 bp or 100 bp to 200 bp and onward.

### 3.2 Virtual evidence’s performance improved regardless of choice of expression threshold

There exists no comprehensive, fully experimentally verified ground truth for TSSs active in some cell type. This is because there exists no universally agreed-upon definition of what it means for a TSS to be active. One possible definition identifies TSS activity based on RNA-seq or global run-on sequencing (GRO-seq)^61^ expression, which measure the gene expression levels of RNA transcripts.^62^ Another definition identifies TSS activity based on histone modification expression at the gene locus.^63,64^ For our analyses, we considered a TSS active in some cell type if it has high CAGE expression in that cell type.^65^

By defining a TSS as active if it has high CAGE expression, we still had to decide on a threshold between “high” and “low” expression.^66^ We could not simply split a set of CAGE tag pileups based on zero or non-zero expression, as regions of low (but non-zero) expression may embody outliers not representative of the bulk sequencing population. This makes the ideal splitting threshold unclear.

To alleviate the need to choose a single threshold for high CAGE expression, we evaluated Segway’s performance on multiple differently-thresholded ground truths. Specifically, we varied the expression threshold used to create the ground truth, and reported Segway’s performance when scored on each thresholded set (Figure 6). This procedure resulted in a curve that effectively reverses the classical precision-recall (PR) curve. To create a PR curve, one varies the classification threshold of a classifier, with fixed ground truth, and plots the precision and recall corresponding to each threshold. Here, instead, we varied the “classification threshold” of the ground truth (the expression threshold required to call an example positive), with fixed classifier. We then plotted the precision and recall corresponding to each threshold. Consequently, we measured the distance of each performance curve to the point (1,1), representing perfect positionally tolerant precision and recall. Over a wide range of expression thresholds, virtual evidence’s performance consistently exceeded that of the unsupervised control (Figure 6).

**Figure 6:**
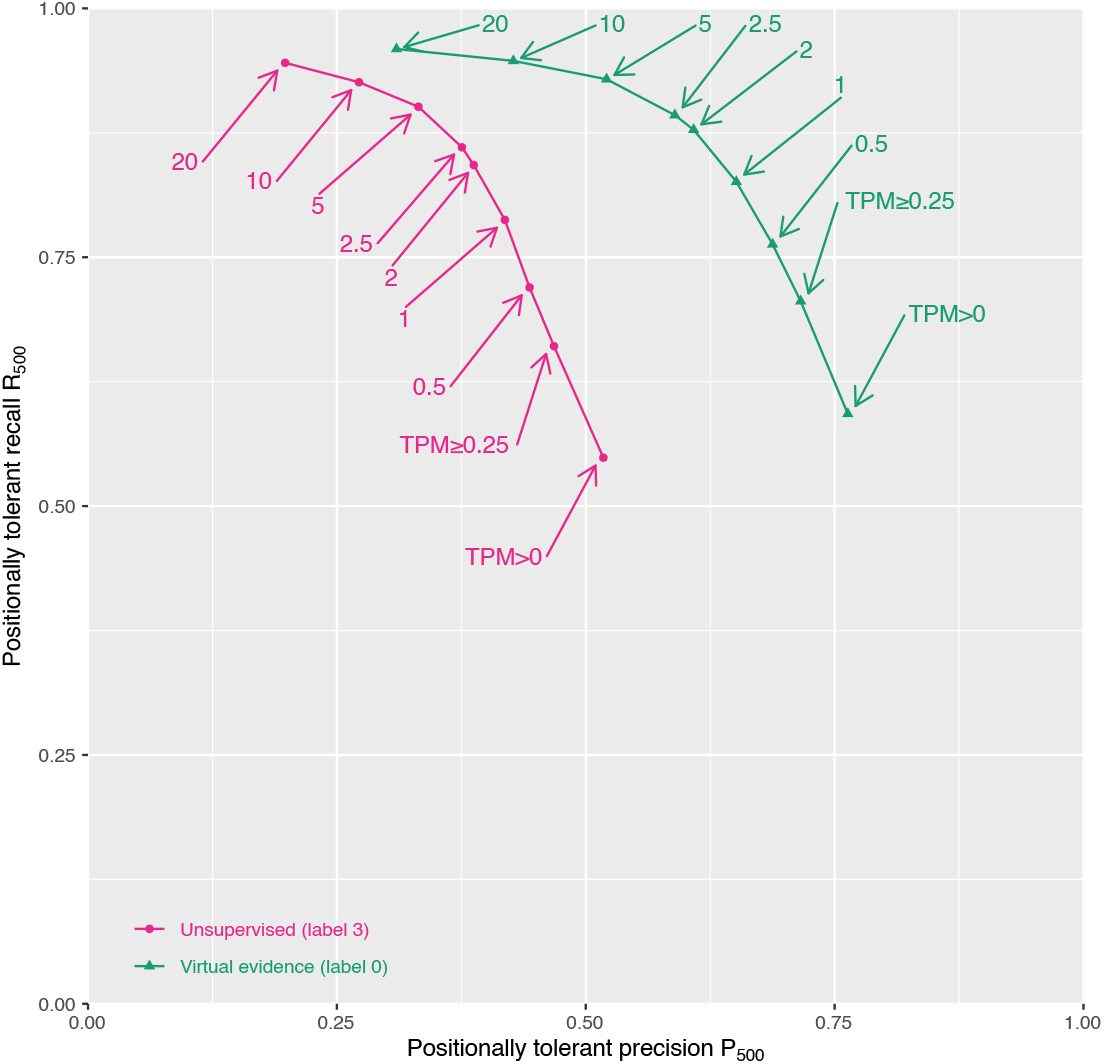
Reverse-PR curve computed with respect to positionally tolerant precision and recall, with a radius of 500 bp. Each curve represents Segway’s performance scored on differently thresholded truth sets. Arrows indicate TPM thresholds. Higher proximity of each curve to the point (1,1) indicates better performance. Pink circles: unsupervised Segway; green triangles: Segway with virtual evidence.

### 3.3 Virtual evidence’s performance remained stable under varying prior equation hyperparameters

To understand the impact of hyperparameter choice on the prior equation (Eqn 4), we trained Segway with varying base and relative confidences and reported each combination’s performance (Table S3). Specifically, we varied the base confidence (from 0.85 to 1.50) and relative confidence (from 0.004 bp/TPM to 0.2 bp/TPM) in the prior equation. We then recomputed the prior probabilities, and re-ran and re-evaluated Segway for each hyperparameter variation. We used a positionally tolerant radius of 500 bp and evaluated on our ground truth of K562 TSS features with TPM ≥ 0.5 located on all chromosomes except chromosome 21. The virtual evidence model’s performance remained stable even with varying prior equation parameters.

## 4 Discussion

Using virtual evidence, Segway learns from provided information on the function of particular regions. Our implementation’s flexibility opens the door to new applications, such as incorporating partial annotations on multiple classes of regions by specifying prior beliefs on several labels simultaneously. For problems where we have no comprehensive, high-confidence ground truth, this ability may improve identifying regions for further biological investigation.

### 4.1 Related work and differences

One can compare Segway with virtual evidence to ModHMM,^67^ a supra-Bayesian^68^ SAGA method which attempts to overcome some of the limitations of unsupervised SAGA methods. As a supra-Bayesian model, ModHMM uses a hidden Markov model, trained on the outputs of multiple heuristic classifiers, to annotate the genome with chromatin states. This provides an approach quite different from Segway with virtual evidence. Segway with virtual evidence incorporates supervision data directly into Segway’s graphical model through dummy virtual evidence nodes. By using only the results of heuristic classifiers to annotate the chromatin state of a region, ModHMM’s supra-Bayesian approach might not capture all nuances of the input data. Moreover, ModHMM trains its hidden states with a fixed transition matrix structure. This necessitates establishing relationships between pre-defined desired chromatin states. Similarly, ModHMM does not train its classifiers on real data, but instead defined them based on heuristics regarding input feature patterns at or around regulatory elements. Also, ModHMM requires input data from a fixed set of assays and histone modifications. Segway and other standard SAGA methods offer far more flexibility and generality of application.

Like Segway with virtual evidence, DECRES^69^ uses a supervised model to identify promoters genome-wide. DECRES, however, uses a multi-layer perceptron. This makes interpreting the trained model more challenging than the generative probabilistic model of Segway.^70^

Sethi et al. [71] supervise on STARR-seq,^72^ DNase-seq, and histone modification data to predict promoters and enhancers across the genome. They do this by combining a shape-based approach with a linear support vector machine. This approach does not generalize, as one cannot use it to perform non-binary classification. In comparison, Segway annotates the genome with a user-defined number of chromatin states, which can be any integer greater or equal to 2.

### 4.2 Future work

Given semi-supervised Segway’s success at predicting promoters, future work could explore its ability to predict rarer and more difficult-to-find elements, such as insulators and enhancers.^56,73^ Future work could also explore the use of other probability functions to compute priors. For example, one could model the noise distribution of CAGE tags and use the probability of non-spurious tags as the prior for TSS. While the tanh function squashes input values between 0 and 1, its output also tends slowly towards 1 given increasing input. This may be undesirable in some situations—for example, when high values require more differentiation in probability. In these cases, one might wish to use a function which grows linearly or faster. Other work could explore the use of multiple supervision labels simultaneously. For example, one could use virtual evidence to supervise on multiple types of gene regulatory regions at once.

## 5 Availability

Segway with virtual evidence (version > 3.0) is available at https://segway.hoffmanlab.org. The version of the Segway source with which we ran our experiments is available at https://doi.org/10.5281/zenodo.3630670. Our code (pre-processing and evaluation scripts), and experimental outputs are available at https://doi.org/10.5281/zenodo.3627261.

## 6 Acknowledgments

We thank Jeffrey A. Bilmes (University of Washington) for his contributions to this work. We thank Carl Virtanen and Zhibin Lu (University Health Network High-Performance Computing Centre and Bioinformatics Core) for technical assistance and Quaid Morris for comments on the manuscript. This work was supported by the Natural Sciences and Engineering Research Council of Canada, the Canadian Institutes of Health Research, the Ontario Ministry of Research, Innovation and Science, and the Princess Margaret Cancer Foundation.

## 7 Supplementary Material

**Table S1:** Datasets used for TSS prediction. All datasets describe the K562 cell line and use the GRCh38/hg38 assembly. All datasets are available at https://www.encodeproject.org/experiments/. Replicates are biological.

| Assay | ENCODE accession ID | Processing | Replicates |
| --- | --- | --- | --- |
| ChIP-seq: H3K27ac | ENCFF779QTH | fold change over control | 2 |
| ChIP-seq: H3K4me3 | ENCFF712XRE | fold change over control | 2 |
| ChIP-seq: H3K36me3 | ENCFF440XMD | fold change over control | 2 |
| ChIP-seq: H3K27me3 | ENCFF928NWQ | fold change over control | 2 |
| DNase-seq | ENCFF868NHV | read-depth normalized signal | 1 |

**Table S2:**
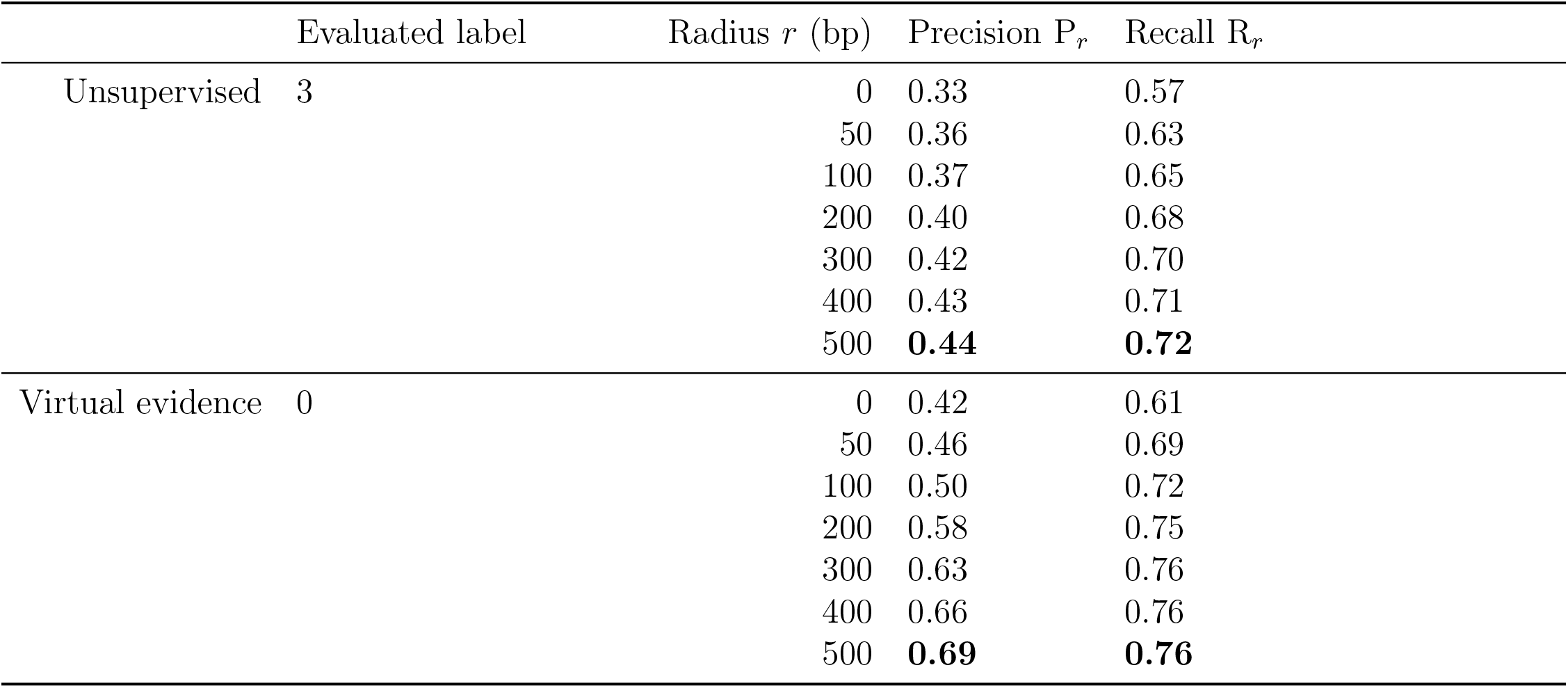
Comparison of performance of virtual evidence against its unsupervised control. Positionally tolerant precision and recall computed with varying radii on the human genome except chromosome 21. Bold: positionally tolerant precision and recall with the radius *r* = 500 bp used elsewhere in this work.

**Table S3:**
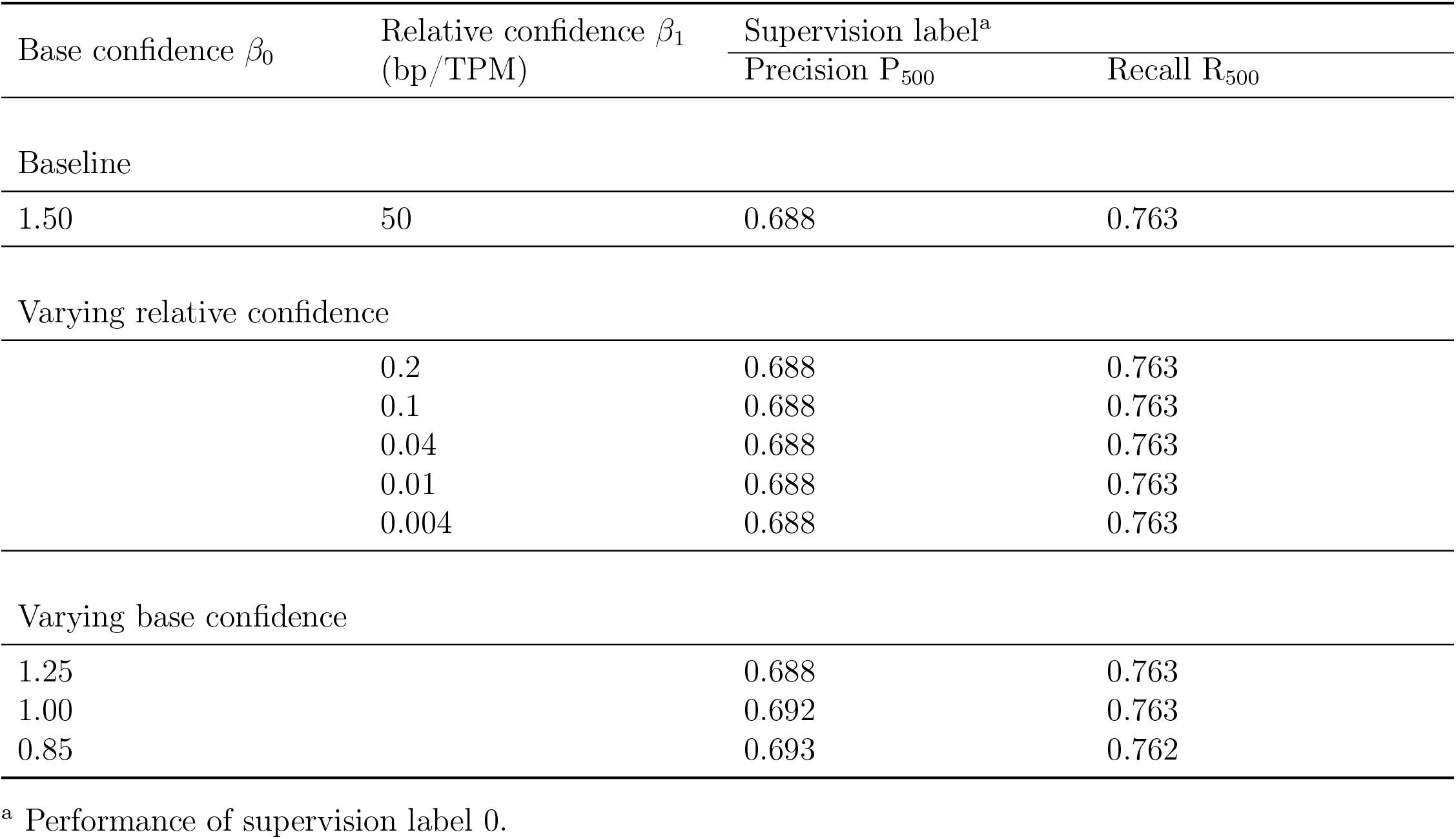
Performance of Segway with virtual evidence under varied prior equation hyperparameters. Positionally tolerant precision and recall computed with the radius *r* = 500 bp. We evaluated each case on our ground truth of K562 TSS features with TPM ≥ 0.5 located on all chromosomes except chromosome 21.

## Notes

https://segway.hoffmanlab.org

https://doi.org/10.5281/zenodo.3630670

https://doi.org/10.5281/zenodo.3627261

